# Olfactory proteomics reveals the capacity of the HDAC1 inhibitor pyroxamide to halt the α-synuclein preformed fibrils-induced damage in nasal epithelial, microglial and dopaminergic neuronal cell lines

**DOI:** 10.1101/2025.10.02.679944

**Authors:** Paz Cartas-Cejudo, Marina De Miguel, Leire Extramiana, Mercedes Lachén-Montes, Silvia Romero-Murillo, Djordje Gveric, Joaquín Fernández-Irigoyen, Enrique Santamaria

## Abstract

Parkinson’s disease (PD) is the second most common neurodegenerative disorder mainly characterized by the degeneration of dopaminergic neurons originating in the substantia nigra (SN) *pars compacta* and projecting to other brain regions, giving rise to motor and non-motor symptoms. Despite significant progress in understanding the molecular and cellular disruptions associated with PD, there remains an unmet clinical need for effective therapies. In this study, proteomic analysis of the olfactory tract (OT) in controls with no known neurological history (n=17) and PD subjects (n=21) revealed Lewy body disease (LBD) stage-dependent proteostatic impairment, accompanied by progressive modulation of the alpha-synuclein (α-syn) functional interactome. Differential OT omic profiles (OMS) were used in a computational drug repurposing approach, reveling the HDAC1 inhibitor pyroxamide as one of the top drug candidates with *in silico* potential to restore altered OMS. To explore the potential therapeutic effects of pyroxamide, *in vitro* assays were performed using α-syn preformed fibrils (PFFs). Pyroxamide treatment reduced α-syn PFFs-induced toxicity in olfactory epithelial, microglial and dopaminergic neuronal cell lines, producing a protective effect against hydrogen peroxide-induced damage exclusively in brain-derived cell types. These findings confirm the suitability of omics profiles in drug repurposing workflows against PD, offering valuable insights into the potential of HDAC1 inhibitors in the therapeutic pipeline of PD.

**Graphical abstract:** 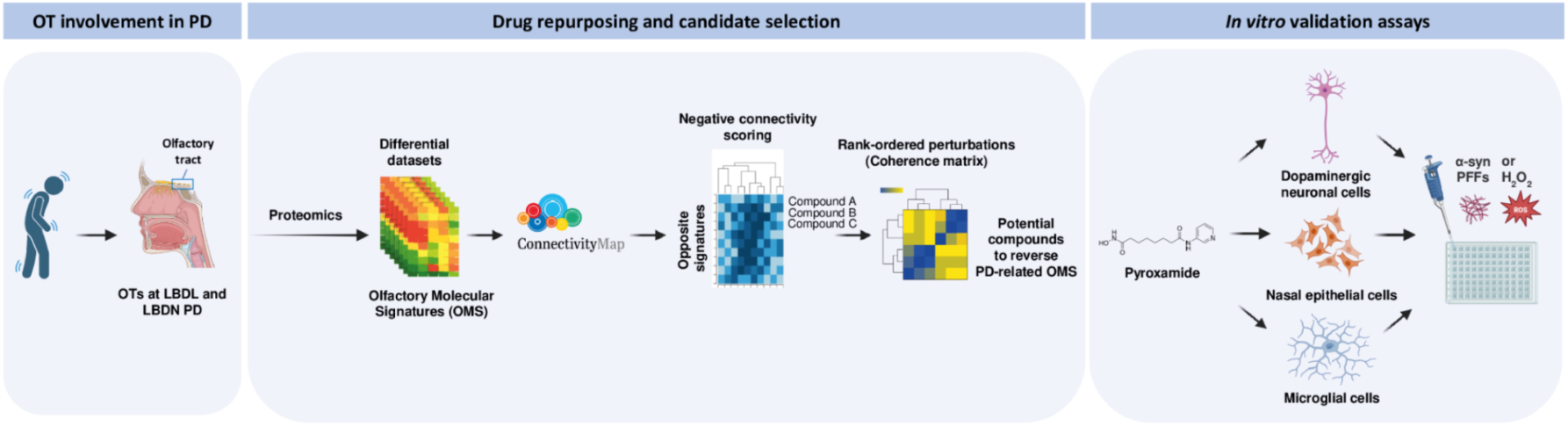

## INTRODUCTION

Parkinson’s disease (PD) is the second most common neurodegenerative disorder [1–3], mainly characterized by the degeneration of dopaminergic neurons originating in the SN *pars compacta* and projecting to other brain regions, marked with motor symptoms, such as tremor, rigidity, and bradykinesia, and non-motor dysfunction, including olfactory besides a typical neuropathological appearance [4,5]. The major pathological features of PD are the deposition of α-synuclein (α-syn) protein in the form of Lewy bodies and Lewy neurites in neurons [6,7]. These α-syn aggregates propagate during disease progression across specific brain regions in a predictable pattern [8], enabling a neuropathologic classification into different Lewy Body Disease (LBD) stages [9,10]. Despite significant progress in understanding PD-related molecular and cellular mechanisms, early disease-related processes remain poorly understood [11]. There is an unmet clinical need to identify new biomarkers for early and presymptomatic diagnosis, as well as for evaluating novel therapies [12]. Besides motor symptoms, olfactory dysfunction occurs in as many as 90% of patients with PD and appears early in the disease course, being suggested as a premotor sign of neurodegeneration [13–15]. Neuropathological changes as well as molecular alterations in olfactory-related areas have been consistently detected in human PD brains [16]. Such pathology in olfactory bulb (OB) and OT could be associated with the severity of PD elsewhere in the brain [17,18]. Axonal degeneration is observed in the OTs of PD subjects, and the loss of the OT fiber integrity correlates with decreased glucose metabolism in central olfactory structures [19,20]. A possible contributing factor to these early alterations is persistent inflammatory activity in peripheral and sensory pathways. Chronic inflammation in the OT and/or the enteric nervous system, potentially triggered by lifelong environmental exposures, pathogens, or gut microbiota imbalances, may promote α-syn aggregation and facilitate its spread to the brain, further aggravating neuroinflammatory responses that contribute to neurodegeneration [21]. Since inflammation is also considered an early event in PD, targeting inflammatory pathways might be more effective in preventing disease progression than in reversing established pathology [22–24]. Therefore, efficient therapies are dependent on the improvement of early diagnosis. One significant strategy toward the delineation of metabolic alterations linked to PD is the generation of molecular expression profiles during disease course across early affected brain regions [25]. A proteome-wide investigation using high-resolution mass spectrometry represents a straightforward strategy for the characterization of the proteome in olfactory regions under diverse biological conditions [26–28]. This unbiased approach has vastly improved the biochemical understanding of PD. However, only a handful of studies have extensively examined the progressive proteostatic imbalance in early-affected areas with the aim of revealing incipient neurodegenerative changes in PD phenotypes [29,30]. We suggest that molecular characterization of the OT’s progressive neurodegeneration in an LBD stage-dependent way will provide a holistic view of the biochemical pathways involved in the olfactory pathophysiology of PD [31]. This will provide the basic knowledge that will enable the computational repurposing of drug candidates that could reverse PD-associated OMS [32,33]. In this work, we integrate neuropathological diagnosis, quantitative proteomics, physical/functional interaction data, large-scale pharmacogenomic databases and *in vitro* functional experiments to understand the chronological regulation of molecular pathways in the OT during the progression of PD as well as to identify and validate repurposed drugs with therapeutic potential against this disease.

## MATERIALS AND METHODS

### Human samples

In compliance with Spanish Law 14/2007 on Biomedical Research, informed written consent for research purposes was obtained from the relatives of patients included in this study through several Spanish Neurological Tissue Banks (Hospital Clínic–Institut d’Investigacions Biomèdiques August Pi i Sunyer-IDIBAPS, IDIBELL Biobank, and Navarrabiomed Biobank). Olfactory tract (OT) specimens (Table 1), and the corresponding clinical and neuropathological data from patients with Parkinson’s disease (PD) were obtained from the Parkinson’s UK Brain Bank (Imperial College London) and from the Neurological Tissue Bank of Navarrabiomed (Pamplona, Spain). Neuropathological assessments were performed in accordance with standardized scoring and grading criteria [34]. In total, twenty-one PD cases were distributed across Lewy body disease (LBD) stages: LBD-limbic (LBDL; n = 11) and LBD-neocortical (LBDN; n = 10). Additionally, seventeen cases from elderly individuals without clinical history or histopathological evidence of neurological disease were included as controls. The study adhered to the principles of the Declaration of Helsinki, and all procedures, including postmortem assessments and experiments, received prior approval from the Clinical Ethics Committee of Navarra Health Service (Study code: PI_2019/108). As previously described, one brain hemisphere was cut into 1-cm thick coronal slices, with the OT region and selected brain areas quickly dissected, frozen on metal plates over dry ice, and stored in individual airtight bags at -80°C for biochemical analysis. The other hemisphere was immersed in 4% buffered formalin for three weeks for morphological examination. Portions of the spinal cord were either frozen at -80°C or fixed in formalin for subsequent analysis. Neuropathological evaluations were performed on paraffin-embedded sections from 25 brain regions, including the cerebrum, cerebellum, brainstem, and spinal cord, stained with hematoxylin and eosin, Klüver-Barrera, and periodic acid Schiff, or subjected to immunohistochemistry using antibodies against β- amyloid, phospho-tau (AT8), α-synuclein, αB-crystallin, TDP-43, TDP-43-P, ubiquitin, p62, glial fibrillary acidic protein, CD68, and IBA1. Neuropathological diagnoses followed current guidelines. Cases with concurrent proteinopathies (such as TDP-43 pathology) were excluded from the study. Control subjects had no history of neurological or mental disorders, and their neuropathological assessment revealed no abnormalities, except for small vessel disease in some cases. A total of 38 postmortem human OT samples were used for proteomic analysis, comprising 17 cases from patients with no history or histological findings of any neurological disease were employed as a control group (n=6F/11M; mean age ± SD: 61.7 ± 10.5 years) and 21 Parkinson’s disease (PD) cases (n=7F/16M; mean age ± SD: 76.9 ± 12 years), distributed across Lewy-type alpha-synucleinopathies (LTS) initial and advanced stages, respectively: LBD-limbic (LBDL; n=11) and LBD-neocortical (LBDN; n=10) (Table 1).

**Table 1:** Description of the OT samples included in this study. Abbreviation: PMI, postmortem interval. M, men. W, women.

|  | Cases | Sex | Age | PMI | Cause of death |
| --- | --- | --- | --- | --- | --- |
| Controls | A1423 | W | 54 | 8h | - |
|  | A1218 | W | 65 | 3h 45m | Agrofilic grain disease |
|  | A1403 | W | 51 | 4h | No neuropathological lesions |
|  | A1576 | W | 60 | 12h | Status cribosus |
|  | A9/43 | W | 51 | 5h 30m | Hypoxia |
|  | A15/7 | W | 54 | 13h 40m | Ischemia, focal |
|  | C048 | M | 68 | 10h | Mild vascular pathology |
|  | C064 | M | 87 | 21h | Cerebellar infarcts, BG vascular pathology |
|  | A1404 | M | 65 | 9h 35m | - |
|  | A1419 | M | 56 | 8h 50m | - |
|  | A1412 | M | 77 | 6h 10m | - |
|  | BCN473 | M | 65 | 3h 05m | - |
|  | A1277 | M | 74 | 9h 25m | Infarction lacunar |
|  | A1368 | M | 45 | 18h 30m | Status cribosus |
|  | A1438 | M | 67 | 5h 50m | Amyloid pathology |
|  | A1377 | M | 59 | 8h 30m | - |
|  | A15/30 | M | 51 | 10h | Tumor, metastasis melanoma |
|  | Cases | Sex | Age | PMI | Stage |
| PD | A15/19 | W | 79 | 13h | LBDI |
|  | A1318 | W | 61 | 18h 30m | LBDII |
|  | A1363 | M | 61 | 13h 20m | LBDII |
|  | BCN496 | W | 74 | 16h | LBDL |
|  | BCN541 | W | 73 | 10h | LBDL |
|  | PD295 | M | 83 | 26h | LBDL |
|  | PD541 | M | 72 | 11h | LBDL |
|  | PD579 | M | 76 | 9h | LBDL |
|  | PD591 | M | 77 | 17h | LBDL |
|  | BCN461 | M | 72 | 9h 10m | LBDL |
|  | BCN525 | M | 73 | 5h | LBDL |
|  | PD495 | W | 88 | 25h | LBDN |
|  | PD501 | W | 89 | 16h | LBDN |
|  | PD550 | W | 83 | 24h | LBDN |
|  | PD357 | M | 71 | 15h | LBDN |
|  | PD450 | M | 66 | 15h | LBDN |
|  | PD537 | M | 84 | 18h | LBDN |
|  | PD636 | M | 84 | 22h | LBDN |
|  | BCN436 | M | 86 | 5h | LBDN |
|  | BCN536 | M | 80 | 21h 35m | LBDN |
|  | CSF-1224 | M | 82 | 7h 25m | LB 4-5 Braak + AgD III + ARP IV B |

### Olfactory proteomics

Protein extraction, proteome quantification, and analysis were performed as previously detailed [30,35], using sequential window acquisition of all theoretical fragment ion spectra mass spectrometry (SWATH-MS) and MS/MS library creation. Human OT samples were lysed in buffer containing 7 M urea, 2 M thiourea, and 50 mM DTT, followed by centrifugation at 100,000 g for one hour at 15°C. Protein concentrations in the supernatants were measured using the Bradford assay (BioRad). To maximize proteome coverage, both in-solution and in-gel digestion methods were employed. For in-solution digestion, the pellet was dissolved in 6 M urea, 100 mM Tris, pH 7.8, reduced with DTT (10 mM) for one hour at 25°C, and alkylated with iodoacetamide (30 mM) for one hour in darkness. An additional reduction step with DTT (30 mM) was carried out at 25°C for one hour. The sample was diluted to 0.6 M urea using Milli-Q water, followed by trypsin digestion (Promega; 1:50, w/w) at 37°C for 16 hours. Digestion was halted by adding acetic acid. Samples were dried by vacuum centrifugation and resuspended in 10 μL of 2% acetonitrile, 0.1% formic acid, and 98% Milli-Q water. To generate the MS/MS library, protein extracts from all samples (1 μg/sample) were pooled, separated by SDS-PAGE (4%-15% stain-free, BioRad), and stained with Coomassie Blue. Thirteen gel slices were excised and enzymatically digested with trypsin (Promega; 1:20, w/w) at 37°C for 16 hours. Peptides were dried, purified using C18 ZipTips (Millipore), and reconstituted to a final concentration of 0.5 μg/μL in 2% acetonitrile, 0.5% formic acid, and 97.5% Milli-Q water for mass spectrometry analysis. MS/MS data were acquired using a TripleTOF 5600+ mass spectrometer (Sciex) coupled with a 2D nanoLC system (SCIEX, Canada). Peptides were separated on a Thermo Scientific C18 column (0.075 x 250 mm, 3 μm particle size, 100 Å pore size) with a gradient from 2% to 40% acetonitrile in 120 minutes at a flow rate of 300 nL/min. The top 35 most intense ions were selected for fragmentation, and spectra were collected from 230 to 1500 m/z. Data acquisition and analysis were performed using ProteinPilot (v5.0, Sciex) with Paragon^TM^ Algorithm for database searches. Proteins were grouped using the ProGroup™ algorithm, and only those identified by at least two unique peptides were considered. False discovery rates (FDR) were calculated using nonlinear fitting [36], with results filtered to 1% global FDR or better. SWATH-MS data were analyzed using PeakView 2.1, with a peptide confidence threshold of 99% (unused score ≥1.3) and a protein FDR lower than 1%. Proteins with at least two unique peptides were included in the quantitative analysis.

### Bioinformatics and Statistical Analysis

In relation to proteomics data, quantitative data from analyzed intensities in PeakView® were processed using Perseus software (v1.6.15.0) [37]. After width-adjustment normalization, Welch’s t-test was applied for statistical comparisons between control and PD groups, with significance set at p < 0.05 and 1% FDR. Proteins showing a fold change above 1.3 or below 0.77 were considered differentially expressed. Boxplots were performed with RStudio (v4.3.1). Pathway analysis was conducted using QIAGEN’s Ingenuity Pathway Analysis (IPA), with significance determined by Fisher’s exact test (p ≤ 0.05). Metascape [38] was also utilized to identify biological functions, with settings of minimum overlap (3), minimum enrichment (1.5), and p < 0.01. Drug repurposing was explored using the Connectivity Map (CMap) platform, as previously described [39]. OT protein datasets from initial and advanced PD stages were used as templates in the CMap tool (https://clue.io) with the aim to identify compounds whose transcriptional effects mirrored or opposed the OMS. The quantitative data obtained from the different experimental *in vitro* procedures were analyzed with the GraphPad program (v10.2.1 395) to obtain the statistical and graphical analysis. A t-test was performed for comparisons between control and drug-based treatment conditions, considering statistical significance when the p-value was less than 0.05.

### In vitro Cell Cultures

Immortalized Human Nasal Epithelial Cells (T9243; Applied Biological Materials, SV40) were cultured, as previously described [35] in DMEM (Gibco, 10741574) high glucose, pyruvate and supplemented with 10% fetal bovine serum (FBS) and penicillin-streptomycin 10000 U/mL (Gibco, 15140122). Cells were maintained at 37°C in 5% CO_2_ and seeded at 1,5 × 10^4 cells/well in 96-well plates. Similar conditions were used for the Dopaminergic Neuronal Cell Line (MN9D; Merck Millipore, SCC281), and the Mouse Microglial Cell Line (BV2; Cytion, 305156), seeded at 1 × 10^4 cells/well in 96-well plates. The experimental phase with the three cell lines commenced 24 hours post-seeding.

### α-syn Soluble Species Preparation

The human recombinant A53T mutant α-syn protein pre-formed fibrils (PFFs) (Type 1) were obtained from StressMarq Biosciences Inc. (SPR-326) in 100 μg vials at a concentration of 2 mg/mL in phosphate buffered saline (PBS) at pH 7.4 and stored at - 80°C. Prior to incorporation into the cells, a sonication process at 30-second intervals for 30 minutes was performed. To simulate the process during the treatment with α-syn PFFs, lipofectamine RNAiMAX (ref. 13778075, Invitrogen), was also used.

### Cell Treatment and MTT Assay

Pyroxamide (HY-13216), entinostat (HY-12163) and apicidin (HY-N6735) were purchased from MedChemExpress and prepared in 1 mL of DMSO at 10 mM. MN9D, T9243, and BV2 cells were pre-treated with different concentrations of pyroxamide (5 and 20 nM) for 3 and 6 hours before or 1 hour after the α-syn PFFs addition for 24 hours, depending on the experimental setup. Cell viability was measured using the MTT assay, where MTT solution (0.5 mg/mL) was added for 2 hours, followed by the addition of DMSO. Absorbance at 590 nm was measured, and results were expressed as cell viability percentages relative to untreated control cells. MN9D, T9243, and BV2 cells were treated with pyroxamide in quintuplicates at two different concentrations according to their IC50 and already employed in previous experimental designs [35]. This procedure was performed simultaneously with the addition of DMSO at the same concentrations. Hydrogen peroxide (H_2_O_2_) was added after an hour of pyroxamide treatment. Cell viability was assessed using the MTT assay at 24 hours post-treatment.

## RESULTS

### Olfactory tract proteostatic disruption and functional specificities during PD neuropathological progression

To assess the complexity and progression of neuropathological changes, the proteomic fingerprint of OT was analyzed across different LBD stages using SWATH-MS quantitative proteomics. Out of 1835 quantified proteins, 98 were differentially expressed between controls and PD phenotypes (Supplementary table 1). To investigate alterations across PD stages, we performed a one-way ANOVA including control, LBDL, and LBDN groups. From this analysis we selected 293 proteins that showed significant differences (p < 0.05). For each of these proteins, intensity values were averaged per group to generate a clustering analysis, that revealed protein subsets with specific protein expression profiles across neuropathological grading (Figure 1). Specifically, we observed protein clusters with a significant aberrant expression that followed different trajectories, exclusively modulated in LBDL or LBDN PD stages (Figure 1A). As shown in figure 1, protein expression profiles of each cluster were associated to a specific functional profile. Cluster C1 corresponded to a protein subset with increased expression across stages (mainly involved in carboxylic acid metabolism) whereas cluster C2 was made up of down-regulated protein intermediates related to modulation of chemical synaptic transmission (Figure 1B). Clusters C3 and C4 were constituted by proteins specifically over-expressed (with functional impact on the regulation of SLITs and ROBOs expression) or downregulated (involved in AMP activated protein kinase signaling) in LBDL stage respectively (Figure 1C). Clusters C5 and C6 were associated to proteins with a stable expression across both stages and functionally related to cell surface presentation of NMDA receptors and neurogenesis (Figure 1D). Clusters C7 and C8 were made up of protein components exclusively modulated in LBDN stage. Specifically, C7 was composed by down-regulated proteins mainly involved in VEGF signaling and vesicle-mediated transport whereas most of up-regulated proteins in cluster C8 corresponded to intermediates of carboxylic acid metabolism (Figure 1E).

**Figure 1:**
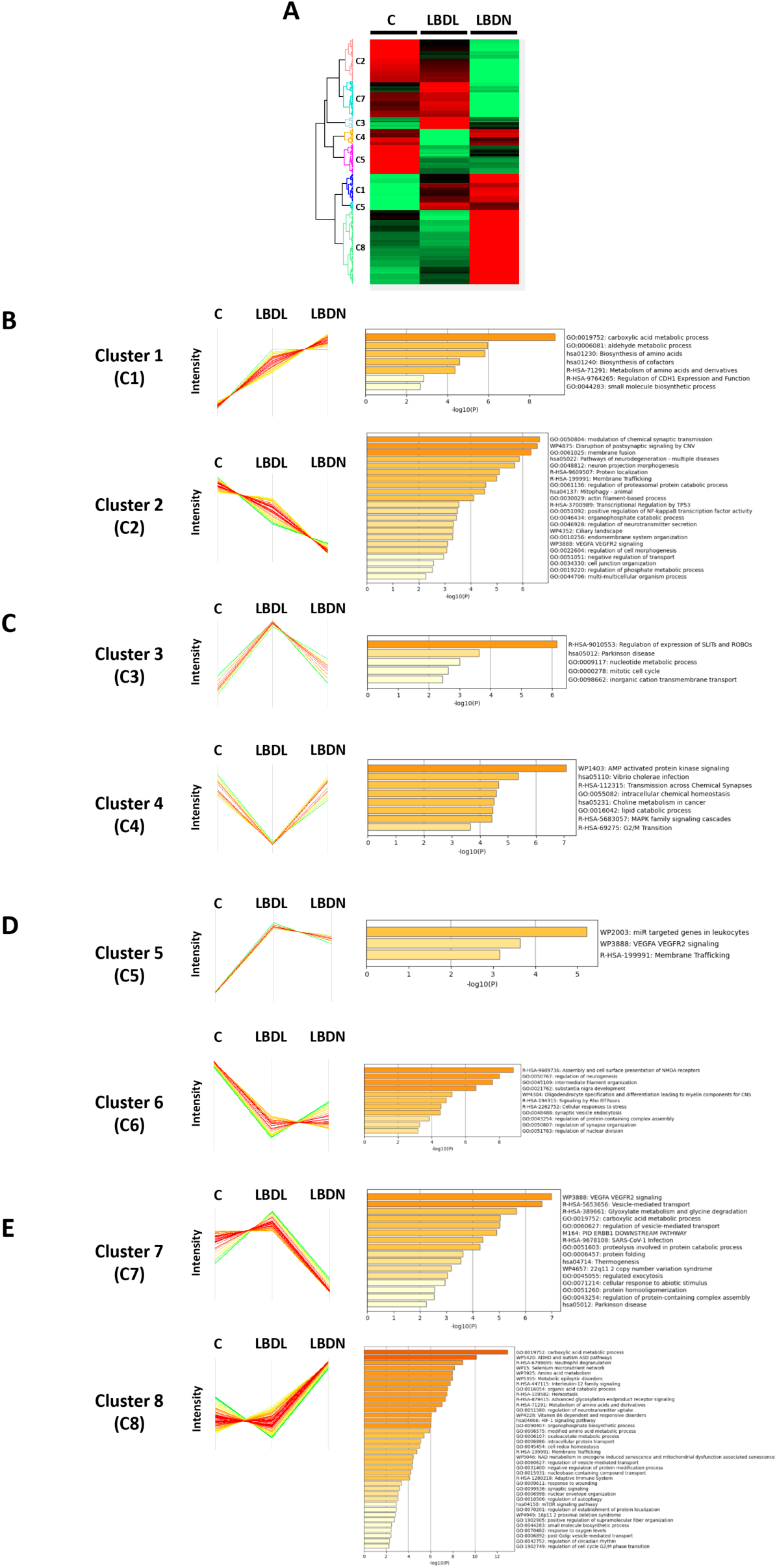
Neuropathological-stage dependent proteomic analysis of the OT in PD. Heatmap representing the differential OT proteotyping in PD across LBD stages (up trend in red; down trend in green) (A). Protein expression and functional profiles associated to each cluster (B-E).

Subsequently, a stage-specific analysis identified 87 and 162 differentially expressed proteins (DEPs) in LBDL and LBDN, respectively (respect to the control group) (Figure 2A- B and supplementary table 1). In LBDL stage, the number of significantly downregulated proteins were slightly higher (56%) in relation to the upregulated ones. In LBDN, practically the number of significant down- or up-regulated proteins was very similar (Figure 2C-D, Supplementary table 1). In addition, twenty-six proteins overlapped between initial and advanced stages mainly involved in intermediate filament organization, substantia nigra development and post-chaperonin tubulin folding pathway (Figure 2E, Supplementary table 1). Moreover, many DEPs across LBD stages were involved in similar biological processes, suggesting functional overlap throughout the different neuropathological stages (Figure 2F). Stage-dependent proteomic datasets were functionally analyzed with the aim of elucidating commonalities and differences between initial and advanced LBD stages. Several common pathways and biofunctions throughout LBD stages share commonalities, highlighting the common alteration during PD olfactory neurodegeneration. However, several processes, such as membrane organization, modulation of chemical synaptic transmission and vesicle-mediated transport are more imbalanced in advances stages. Regarding initial stages, other pathways present more alteration in relation to advanced stages, including nervous system development, intermediate filament organization, post NMDA receptor activation events and actin cytoskeleton organization (Figure 2G). At the subcellular level, common components were differentially affected across LBD staging. As expected, since the OT is primarily composed of axonal bundles derived from the OB, GO terms related to microtubule, actin cytoskeleton, myelin sheath, filopodium, focal adhesion, axon, postsynaptic intermediate filament cytoskeleton and intermediate filament are commonly affected between both PD stages. However, there were some cellular components specifically disrupted in LBDL stage, such as cytoplasmic ribonucleoprotein granule, cytosolic ribosome, glial cell projection, secretory granule membrane, acrosomal vesicle and clathrin-coated pit. On the other hand, multiple altered functionalities were detected in LBDN, focused on presynapse, followed by others including ficolin-1-rich granule lumen, calcium- and calmodulin-dependent protein kinase complex and endocytic vesicle (Figure 2H).

**Figure 2:**
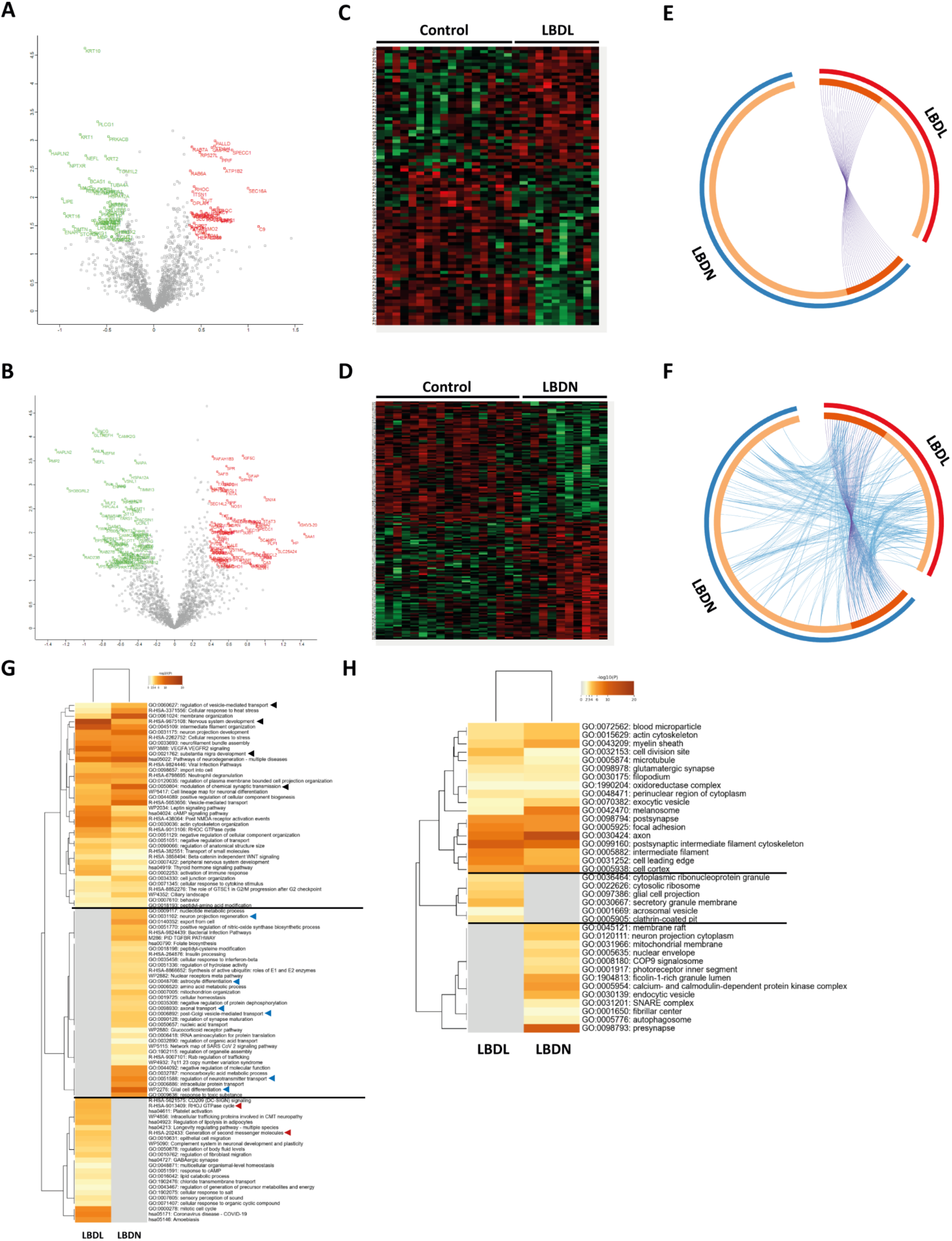
Neuropathological-stage dependent proteomic alterations in the OT and their functional impact in PD stages. Volcano plots reflecting the increasing proteostatic disturbance across LBDL (A) and LBDN stage (B). Heatmap representing the differential OT distribution between controls and LBDL (C), and LBDN (D) (red: up-regulated DEPs; green: down-regulated DEPs). Circos-plot depicting the OT impaired proteome shared across initial and advanced stages (purple lines) (E). Circos-plot showing the functional overlap (blue lines) between deregulated proteomes in both stages at the OT level (F). Functional mapping of the dysregulated OT proteome in LBDL and LBDN (G). Functional mapping of impaired OT proteome across PD grading at subcellular level (H).

### Disruptions in PD-associated protein interactomes during OT neurodegeneration

To determine if impaired OT protein mediators were directly linked to neuropathological protein networks, we performed a systems biology approach that enabled us to partially analyze the alteration of functional protein networks related to α-syn (*SNCA*) metabolism. Considering BIOGRID experimental repository [40], eight proteins were predicted to interact with *SNCA* (*BCAS1*, *CAMK2B*, *INA*, *NEFL*, *NEFM*, *PAK1*, *SEC16A*, *TUBB6*). Intriguingly, all of them were downregulated in both initial and advanced stages, except for SEC16A, that was the only upregulated protein in LBDL and LBDN (Figure 3A-B). In addition, a systems-biology approach allowed us to partially characterize the differential modulation of functional protein networks associated with *SNCA* in initial and advanced LBD stages of PD, revealing multiple deregulated OT proteins as functional *SNCA* interactors (Figures 4A-B).

**Figure 3:**
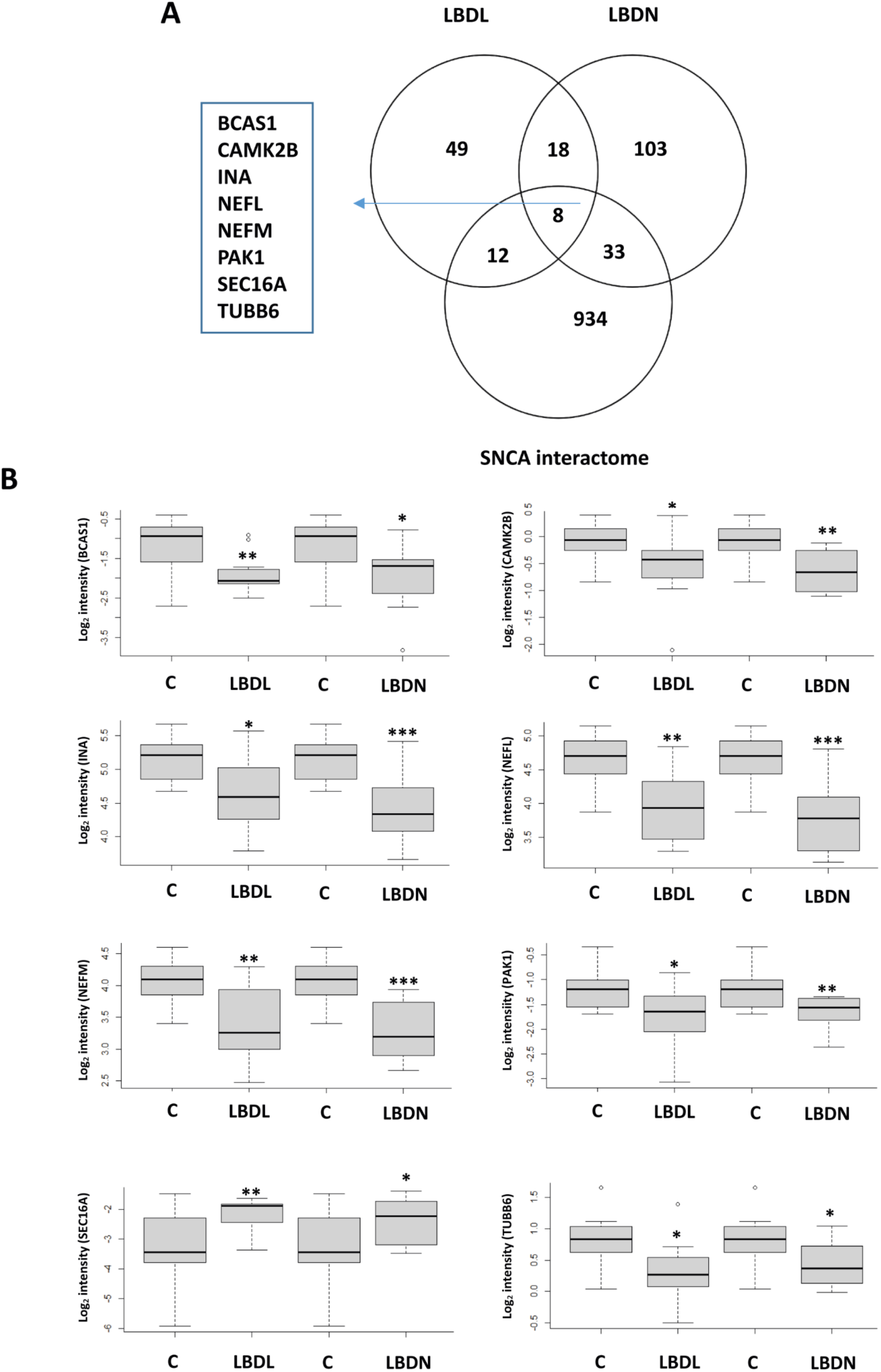
OT-driven alterations within the *SNCA* interactome. Venn diagram representing the overlap between OT-altered proteins and protein components of *SNCA* interactome according to BIOGRID repository (A). Mass spectrometry-based protein intensity of deregulated *SNCA* interactors. Boxplots show width-adjusted normalized protein intensities (B). * indicates p < 0.05; ** p < 0.01; *** p < 0.001.

**Figure 4:**
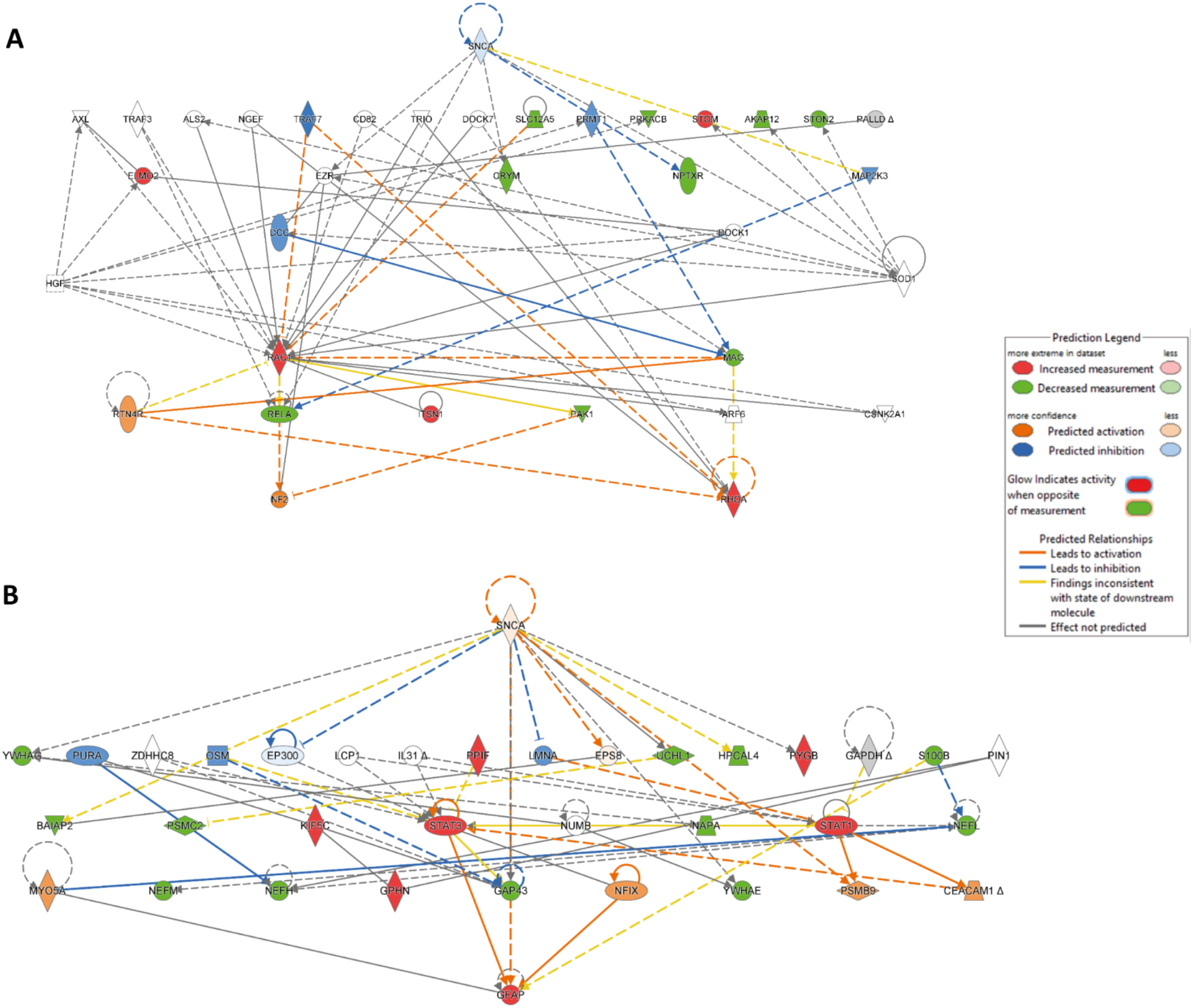
*SNCA* network dysfunction in initial and advanced stages of LBD at the OT level. Deregulation of *SNCA* functional network in LBD initial (A) and advanced (B) stages at the OT level according to IPA knowledgebase.

### Drug repurposing and candidate selection from PD-related omics signatures

Following the drug repurposing workflow previously described in [39], we applied a computational strategy using both OT differential proteomics datasets associated to LBDL and LBDN stages. Using the CMap framework, we selected the most represented candidates across both OMS belonging to the leading pharmaceutical class (HDAC inhibitors) identified in our analysis (connectivity score below –80): pyroxamide, entinostat, and apicidin. This indicates that compounds with highly negative values could have the potential to in silico reverse pathological molecular profiles associated to OT area in PD. First, MTT assays were used to determine the non-cytotoxic concentration ranges for each compound in dopaminergic, olfactory, and microglial cell lines. As shown in supplementary figure 1, none of the compounds showed toxicity at the selected concentrations, except for apicidin in BV2 cell line. After the establishment of *in vitro* α-syn PFFs-induced neurotoxicity assays, different treatment conditions were analyzed. As shown in figure 5A, pyroxamide pre-treatment (20nM) showed neuroprotective effect in dopaminergic MN9D cells when administered 6 hours prior α- syn PFFs exposure. Moreover, the survival potential was also significantly increased when α-syn PFFs were added to the cell culture 1 hour before than pyroxamide (Supplementary figure 2A). A significant survival increment was also observed in olfactory T9243 cells pre-treated with pyroxamide (3 and 6 hours) at 5 and 20 nM, without detecting a dose-dependent effect (Figure 5B). Similar to the observed phenomena in dopaminergic cells, a significant protection effect was observed when α- syn PFFs were administered 1 hour prior to pyroxamide (20nM) (Supplementary figure 2B). Respect to α-syn PFFs-treated microglial BV2 cells, pyroxamide pre-treatment (3h) was able to significantly increment the survival potential at both concentrations (5 and 20nM) (Figure 5C). Upon H₂O₂-induced oxidative stress assays, pyroxamide pre-treatment also restored the survival capacity in dopaminergic MN9D cells (3 and 6h, 20nM) (Figure 6A) as well as in microglial BV2 cells (3h, 20nM) (Figure 6B). No changes were observed in olfactory epithelial cells upon oxidative stress conditions. Neither entinostat nor apicidin exhibited protective effects under any of the conditions or time points tested (data not shown).

**Figure 5:**
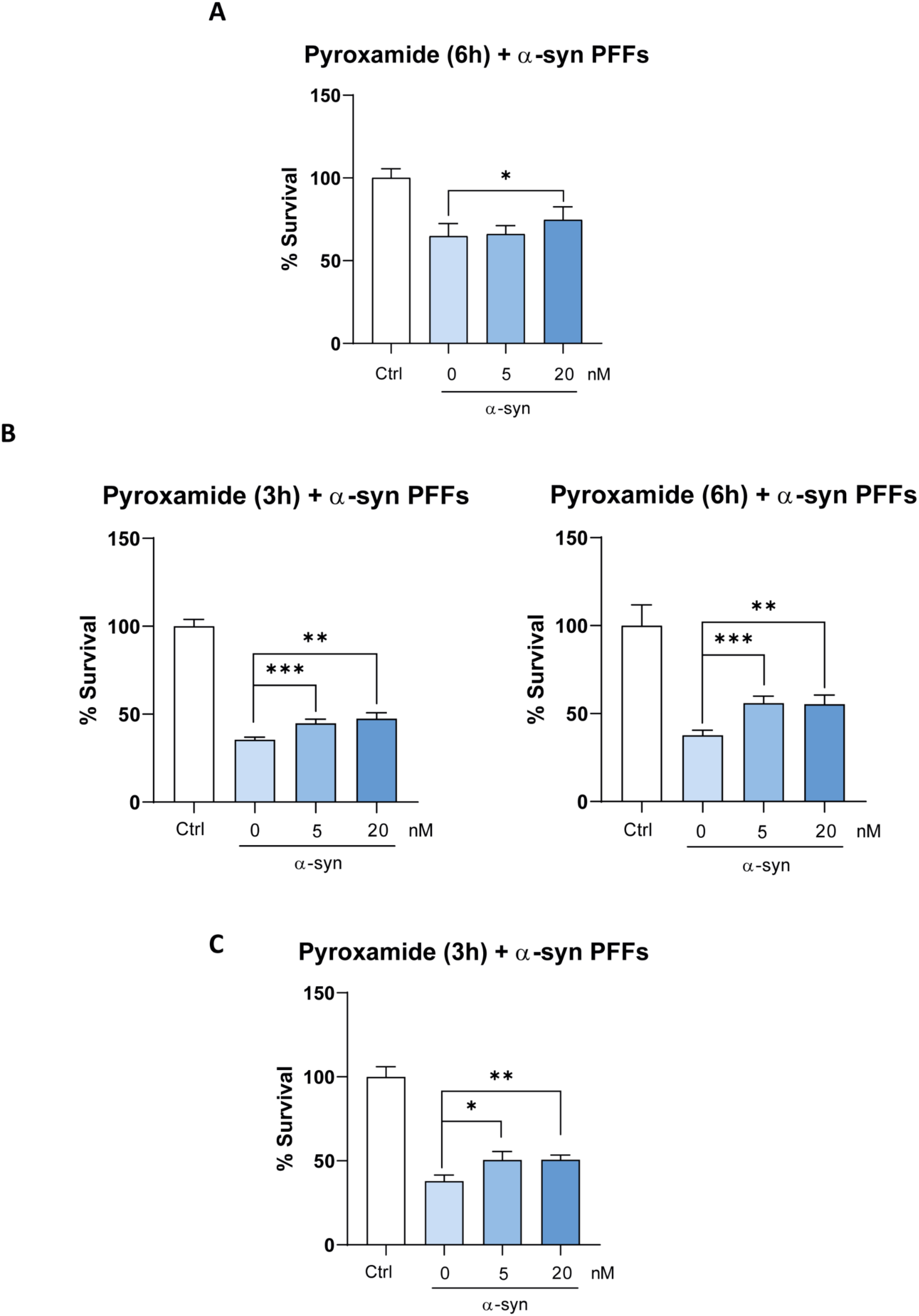
*In vitro* validation assays using pyroxamide upon α-syn PFFs neuropathological insult. *In vitro* experiments in dopaminergic MN9D cells representing the survival effect of α-syn PFFs after 6h of pyroxamide pre-treatment (A). Survival effect of α-syn PFFs after 3 and 6 hours of pyroxamide pre-treatment in olfactory epithelial T9243 cells (B). Survival effect of α-syn PFFs after 3 hours of pyroxamide pre-treatment in microglial BV2 cells upon α-syn PFFs-induced damage (C). Data were analyzed using one-way ANOVA. * indicates p < 0.05; ** p < 0.01; *** p < 0.001.

**Figure 6:**
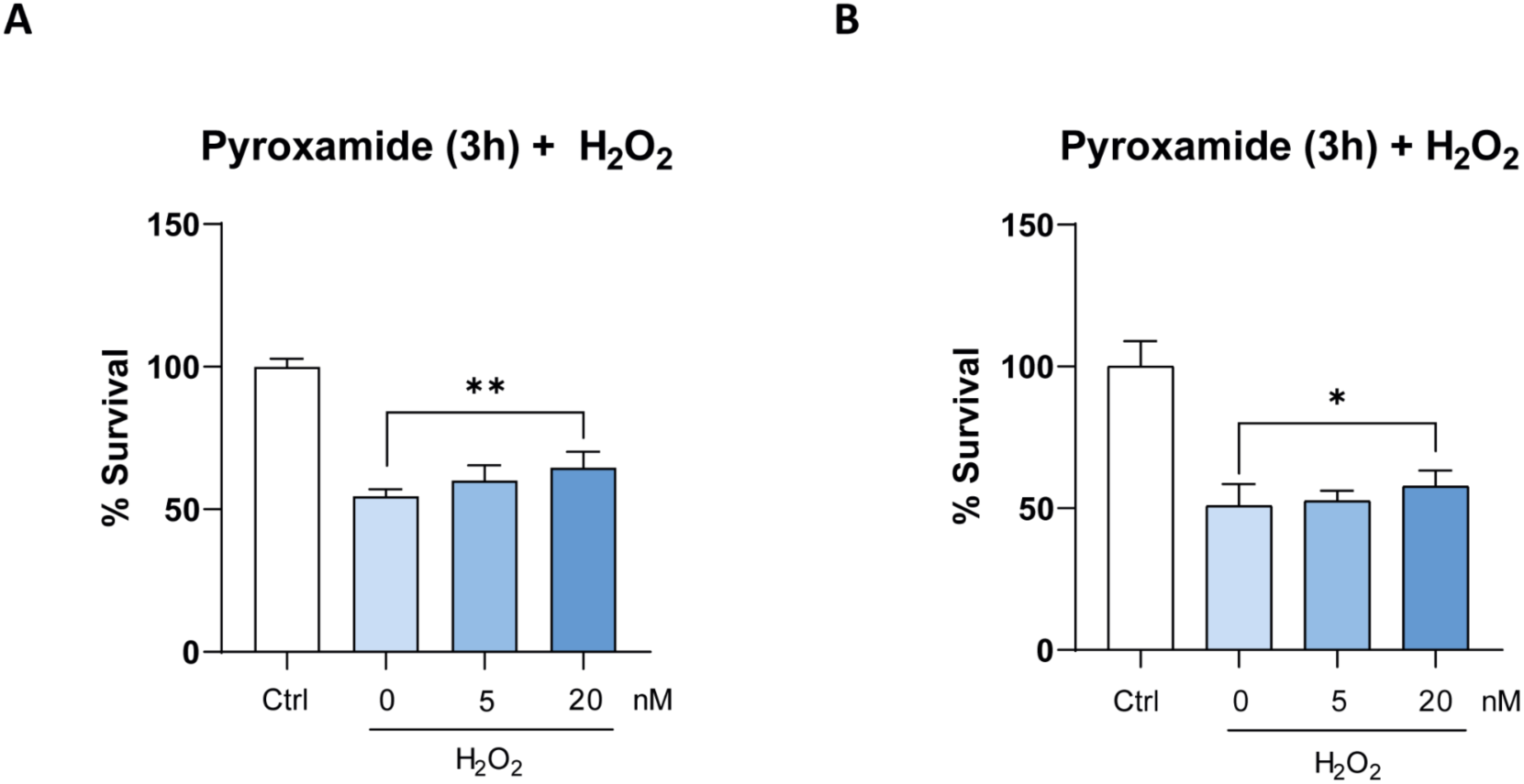
*In vitro* validation assays using pyroxamide upon H_2_O_2-_induced oxidative stress. Survival analysis associated to pyroxamide pre-treatment 3 hours in dopaminergic neuronal (A) and microglial cells (B) upon oxidative stress conditions. Data were analyzed using one-way ANOVA. * indicates p < 0.05; ** p < 0.01.

## DISCUSSION

Parkinson’s disease (PD) is a prevalent and debilitating neurodegenerative disorder with no curative treatment, imposing substantial economic and social burdens. Disease progression is associated with the intensification of comorbidities and a decline in quality of life, nearly doubling patient mortality rates. Current therapeutic strategies are largely symptomatic, failing to halt disease progression, which underscores the urgent need for novel interventions [41]. Non-motor manifestations, including olfactory dysfunction, have gained attention as early indicators of PD [14,15,42], highlighting the importance of understanding initial pathological events for the development of early interventions that may slow neurodegeneration. In this context, we performed quantitative olfactory tract proteomics across both early and advanced PD stages to elucidate the molecular mechanisms underlying olfactory deficits and their temporal progression. While several transcriptomic and proteomic studies have explored PD-affected brain regions [29,43–46], high-throughput molecular insights focused on the olfactory axis remain largely unexplored. One primary objective of this study was to generate comprehensive functional data on protein subsets associated with OT neurodegenerative progression. Our findings demonstrate that the chronological molecular perturbation in the OT diverges significantly from the proteome alterations observed in early disease stages in regions such as the SN, prefrontal cortex, or amygdala [27,43,47], suggesting a lack of coordinated transcriptional programs across brain structures during early olfactory neurodegeneration. Among the differentially expressed proteins identified, eight experimentally validated *SNCA* interactors were dysregulated in the OT, all downregulated in both initial and advanced stages, except for SEC16A, a critical protein regulated at endoplasmic reticulum exit sites by LRRK2, a kinase implicated in PD [48,49]. Upregulation of SEC16A during disease progression could disrupt protein secretion and cellular homeostasis, contributing to neurodegeneration [50,51]. Current PD pharmacotherapy targets motor symptoms [52–56], leaving non-motor symptoms and disease progression unaddressed, which prompted an omic-based drug repurposing approach [35]. Differentially expressed OT proteins from early and advanced stages were analyzed using CMap, revealing three compounds: pyroxamide, entinostat, and apicidin, that potentially reverse PD-associated omic profiles. Pyroxamide selectively inhibits HDAC1, entinostat inhibits HDAC1–3, and apicidin targets HDAC2 and HDAC3, with minor activity against HDAC7 and HDAC8 [57,58]. HDACs are pivotal epigenetic regulators implicated in both cancer and neurodegenerative disorders. Specifically, HDAC1 and HDAC2 are linked to neurodegeneration by impairing axonal transport and reducing mitochondrial activity [59], processes significantly altered in PD OTs. Based on our results, only the HDAC1 inhibitor pyroxamide was able to counteract the survival imbalance induced by α-syn PFFs in olfactory epithelial, dopaminergic neuronal and microglial cells. According to our data, the HDAC1 inhibitor suberoylanilide hydroxamic acid (SAHA), protects dopaminergic neurons from MPP+-induced toxicity, enhancing neurotrophic factor expression and reducing oxidative stress, inflammation, and apoptosis [60]. Similarly, valproic acid improves dopaminergic neuron survival, reduces neuroinflammation, and ameliorates motor deficits in LRRK2 R1441G transgenic PD mice [61], while pharmacological inhibition of HDAC1 (MS-275; entinostat) and HDAC6 (tubastatin) in zebrafish models enhances dopaminergic neuron survival and motor function [62]. Mechanistically, HDAC1 acts synergistically with protein phosphatase 1 gamma (PP1γ) to promote CREB dephosphorylation and inactivation, contributing to dopaminergic degeneration [63]. Selective HDAC1/2 inhibition alleviates motor deficits and neurodegeneration in PD models, reduces neuroinflammation, and improves dopaminergic neuron survival [64]. Pan-HDAC inhibitors also confer neuroprotection, regulating neuronal and glial gene expression, exemplified by cinnamyl sulphonamide hydroxamate derivatives with effects comparable to valproic acid in MPTP-induced rat models [65]. Despite this, the neuroprotective potential of specific HDAC isoforms remains to be fully characterized [62]. Our data-driven drug repurposing approach thus identifies HDAC inhibition as a promising strategy for PD, warranting further investigation of compound synergy in alleviating PD pathology, including potential intranasal administration to target early olfactory deficits, given the predicted limited blood-brain barrier penetration of pyroxamide [66]. Beyond dopaminergic neurons, early PD stages involve olfactory epithelium and microglial alterations [67–73].HDAC inhibitors modulate neuroimmune responses, and protect dopaminergic neurons from LPS- and MPTP-induced toxicity [74,75]. Proteins related to inflammatory pathways, such as RAC1, NF2, STAT1, and STAT3, were differentially regulated in early PD OT datasets, suggesting early activation of inflammatory and stress responses, aligning with previous observations of microglial overactivation contributing to neurodegeneration [76,77]. Additionally, HDAC inhibition enhances expression of neurotrophic factors like GDNF and BDNF in astrocytes [78] and rescues olfactory learning deficits in Drosophila via PQBP1 modulation [79], indicating a broad neuroprotective potential. Taken together, our findings support the hypothesis that HDAC inhibition, including isoform-specific compounds identified in our study, may offer a multipronged therapeutic approach targeting dopaminergic neurons, olfactory deficits, and neuroinflammation. Moving forward, pyroxamide should be evaluated *in vivo* testing in PD models, potentially via intranasal delivery to bypass blood-brain barrier limitations and directly addressing early olfactory dysfunction. Such approaches may pave the way for early intervention strategies capable of modulating disease trajectory and alleviating both motor and non-motor PD manifestations.

## LIMITATIONS

A significant limitation of our study is the difficulty in obtaining postmortem samples with comparable ages between the control and pathological groups, which could introduce potential biases in the interpretation of our results. In addition, the presence of co-pathologies in both the control and pathological sample sets further complicates the analysis, as these co-existing conditions may influence the underlying biological mechanisms and obscure the specific effects related to the primary pathology being investigated. Furthermore, our dataset has limited coverage of low-abundance proteins, hydrophobic proteins, and receptors that are potentially significant in the context of olfactory neurodegeneration. These molecules, which may play a crucial role in the pathophysiology of the disease, are not adequately represented in our findings due to technological constraints inherent to the mass-spectrometry phase. Addressing these limitations, such as obtaining more age-matched postmortem samples, ensuring a more comprehensive inclusion of proteins and receptors, and expanding sample sizes for better statistical power, will be essential for advancing our understanding of the neurodegenerative processes involved in PD at the level of olfactory areas. Finally, we are aware that the drug repurposing workflow used is based on pharmacological profiles obtained in human cancer cell lines. This means that brain specific processes such as synaptic transmission are excluded, indicating that the reversion of neuroanatomical omics profiles may not be perfectly adapted, being mostly represented by genes involved in other type of biological process such as apoptosis, differentiation and inflammation.

## Supporting information

OT quantitative proteomics data and differential OT proteins detected in PD across LBD stages.

## Author contributions

Conceptualization: Enrique Santamaría; sample collection & neuropathology: Djordje Gveric; data curation: Paz Cartas-Cejudo, Marina De Miguel, Mercedes Lachén-Montes, Joaquín Fernández-Irigoyen, and Enrique Santamaría; formal analysis: Paz Cartas-Cejudo, Marina De Miguel, Leire Extramiana, Mercedes Lachén-Montes, Silvia Romero-Murillo, Djordje Gveric, Joaquín Fernández-Irigoyen, Enrique Santamaria; *in vitro* studies: Paz Cartas-Cejudo, Marina De Miguel, Leire Extramiana, Silvia Romero-Murillo; funding acquisition: Enrique Santamaría, Joaquín Fernández-Irigoyen; investigation: Paz Cartas-Cejudo, Marina De Miguel, Leire Extramiana, Mercedes Lachén-Montes, Silvia Romero-Murillo, Djordje Gveric, Joaquín Fernández-Irigoyen, Enrique Santamaria; methodology: Paz Cartas-Cejudo, Marina De Miguel, Leire Extramiana, Mercedes Lachén-Montes, Silvia Romero-Murillo, Joaquín Fernández-Irigoyen, Enrique Santamaria; writing—original draft: Paz Cartas-Cejudo and Enrique Santamaría; and all authors gave final approval of the manuscript and are accountable for all aspects of the work.

## Acknowledgements

We are very grateful to the patients and relatives who generously donor the brain tissue for research purposes. We are indebted to the Parkinson’s UK Brain Bank funded by Parkinson’s UK, a charity registered in England and Wales (258197) and in Scotland (SC037554), the Neurological Tissue Bank of the Biobank from the Hospital Clinic-Institut d’Investigacions Biomédiques August Pi i Sunyer (IDIBAPS, Barcelona), the Neurological Tissue Bank of HUB-ICO-IDIBELL (Barcelona, Spain) and the Neurological Tissue Bank of Navarrabiomed (Pamplona, Spain) for providing us the olfactory specimens as well as the associated clinico-pathological data. Authors thank the PRIDE Team for helping with the mass spectrometric data deposit in ProteomeXChange/PRIDE. The Clinical Neuroproteomics Unit of Navarrabiomed is a member of the Spanish Olfactory Network (ROE) (supported by grant RED2022-134081-T funded by Spanish Ministry of Science and Innovation). The graphical abstract was created using <u>Biorender.com</u>.

## Data availability statement

Mass-spectrometry data and search results files were deposited in the Proteome Xchange Consortium via the JPOST partner repository (https://repository.jpostdb.org) [80] with the identifier PXD068256 for ProteomeXchange and JPST004066 for jPOST (for reviewers: https://repository.jpostdb.org/preview/91792741868c19915ae947 Access key: 3343). According to recent recommendations [81], sex annotation has been included in raw files to facilitate further analysis.

## Ethics statement

According to the Declaration of Helsinki, all assessments, postmortem evaluations, and experimental procedures were previously approved by the Clinical Ethics Committee of Navarra Health Service (study code: PI_2019/108).

## Patient consent statement

According to the Spanish Law 14/2007 of Biomedical Research, informed written consent from several Spanish and British Neurological Tissue Banks was obtained for research purposes from relatives of subjects included in this study.

## Supplementary figures

**Supplementary figure 1:**
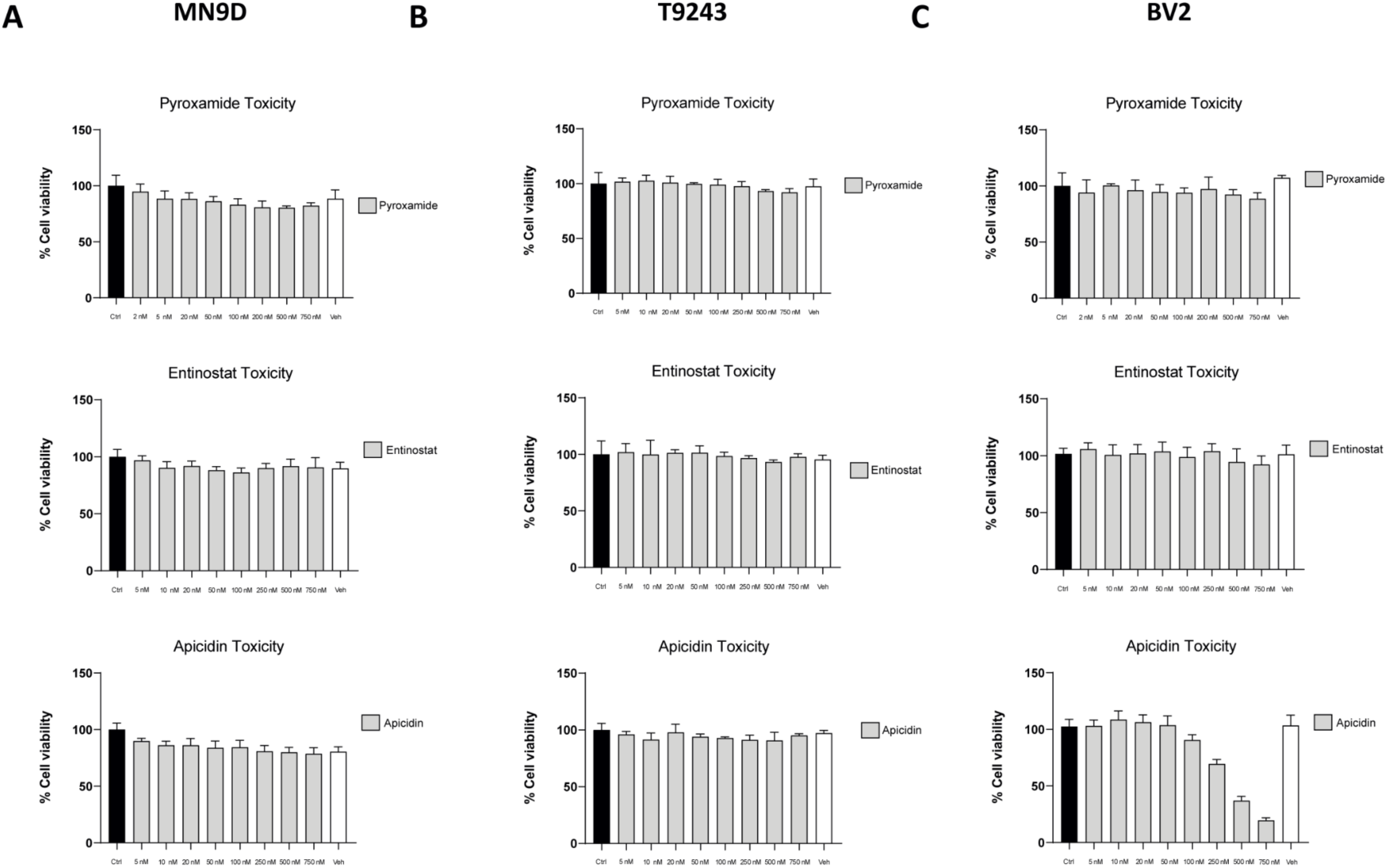
Effect of different concentrations of the selected compounds on cell viability of MN9D, T9243 and BV2 cells. Cells were treated with compounds for 24 h. Results are presented as % of survival and expressed as mean ± SEM (n ≥ 6). MN9D (A), T9243 (B) and BV2 (C) cell viability upon pyroxamide, entinostat and apicidin treatment.

**Supplementary figure 2:**
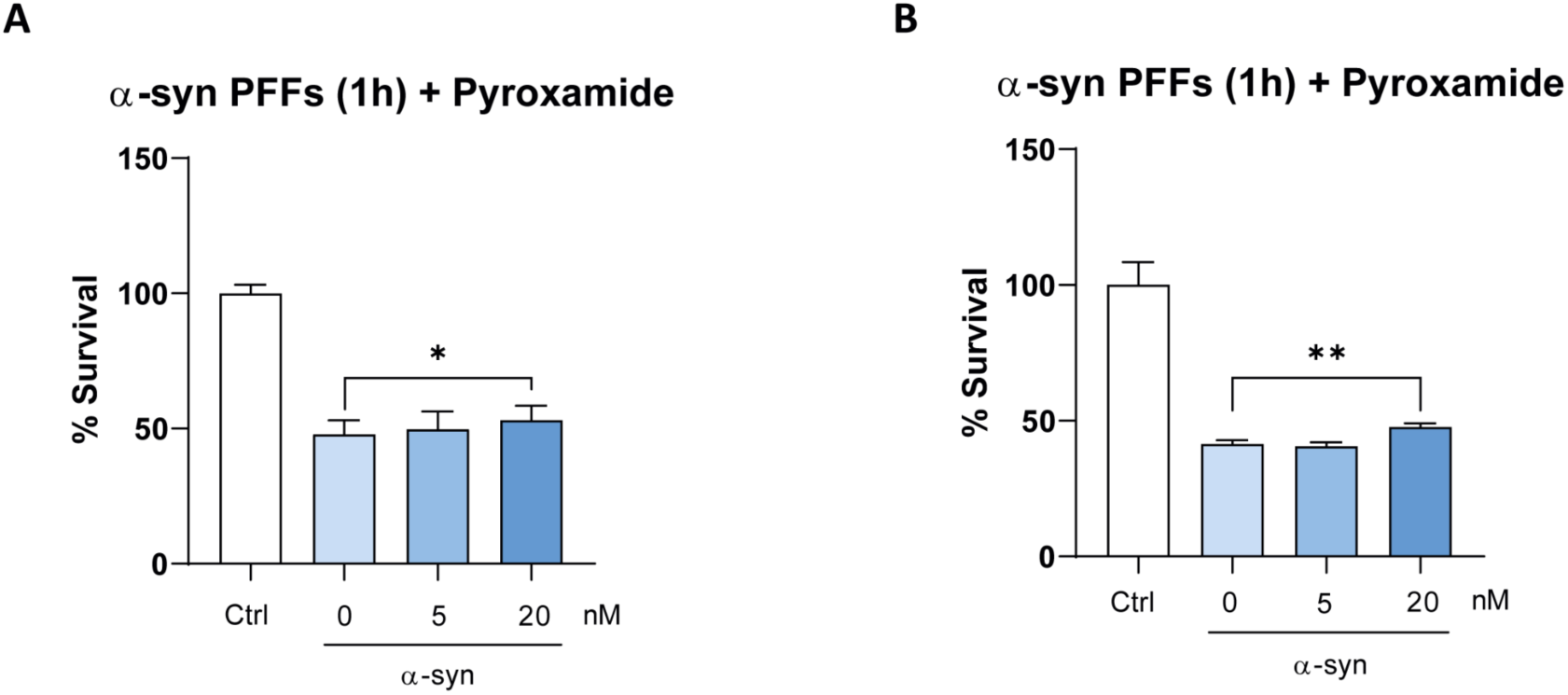
*In vitro* validation assays with pyroxamide upon α-syn PFFs insult. Survival analysis associated to α-syn PFFs administration 1 hour prior to pyroxamide treatment in dopaminergic neuronal (A) and nasal epithelial cells (B). Data were analyzed using one-way ANOVA. * indicates p < 0.05; ** p < 0.01.

**Supplementary table 1: OT quantitative proteomics data and differential OT proteins detected in PD across LBD stages.**

